# Enhancing Collagen Biosynthesis in Mammalian Cells Through Hypoxia-Mimetic Prolyl Hydroxylase Inhibition

**DOI:** 10.1101/2024.12.25.630305

**Authors:** Bar Shahar, Itay Kilimnik, Lucia Adriana Lifshits, Francesca Netti, Marina Sova, Dalia Rosin-Grunewald, Maayan Gal, Lihi Adler-Abramovich

**Affiliations:** Department of Oral Biology, The Goldschleger School of Dental Medicine, Faculty of Medical and Health Sciences; The Jan Koum Center for Nanoscience and Nanotechnology, Tel Aviv University, Israel; The Center for the Physics and Chemistry of Living Systems, Tel Aviv University, Israel; Arrakis Bio LTD, Israel

## Abstract

Collagen, the most abundant protein in the extracellular matrix of mammalian cells, is extensively needed in various biotechnological and therapeutic applications, such as tissue engineering and regeneration, cosmetics, and cultivated meat. Despite the increasing demand for natural collagen from non-animal sources, it is mainly produced from animal connective tissues. Recent research has highlighted that under hypoxia, the activation of the hypoxia-inducible factor (HIF) leads to enhanced collagen type I biosynthesis. However, under normal oxygen conditions, HIF activity is downregulated by the HIF-prolyl hydroxylase (PHD) enzyme. We, therefore, hypothesized that inhibiting PHD could elevate HIF transcriptional activity and enhance collagen biosynthesis under normoxia. Our study demonstrates that inhibiting PHD using exogenous small molecules boosts HIF activity and upregulates the key enzymes, collagen prolyl 4-hydroxylases and lysyl hydroxylases, resulting in up to 29-fold increase in collagen type I in embryonic mouse fibroblast NIH/3T3 cells. These findings suggest that targeting PHD can effectively enhance collagen production in mammalian cells. Therefore, modulating key protein signaling pathways presents a promising strategy for enhancing the production of high-yield natural collagen.

## Introduction

Collagen is a key protein in the extracellular matrix (ECM) of connective tissues of tendons and ligaments, as well as in the skin and bones^1^. Its primary functions are to maintain the structural integrity of the body, regulate cell adhesion, and direct tissue morphogenesis, differentiation, and homeostasis^2^. Collagen type I, the predominant form among 29 distinct types of collagens^3^, is a heterotrimer that assembles into a fibrillar triple-helix structure composed of two α1(I) chains and one α2(I) chain^1^. Its supramolecular structure imparts collagen type I with its unique physiochemical characteristics, including biocompatibility, sustainability, low immunogenicity, and stability. These attributes make it an exceptional bio-scaffold widely used in biotechnological applications such as tissue engineering^4^, pharmaceuticals, cosmetics, and regenerative medicine^5–7^. As collagen scaffolds aspirate to recapitulate the complex native tissue, collagen-based products are highly variable, including tissue grafts, hydrogels, sponges, fibers, films, hollow spheres, and tissue-engineered living substitutes^3^. Specifically, collagen and its derivative gelatin are extensively used in the cultivated meat and food industry^8,9^, supporting the growth of 3D tissue culture in an ECM-mimicking environment^10^.

The primary source of collagen is obtained by extracting it from animal connective tissues^3^. Although this process is cost-effective, major concerns remain regarding immunogenicity and potential disease transmission across species, particularly in medical-related applications^11^. These challenges incentivize research for alternative collagen sources. Indeed, collagen can be produced *in vitro* in cells or recombinantly in expression systems such as plants^12^ and various microorganisms^13^. Moreover, collagen fiber’s structure can be mimicked by synthesizing the minimal canonical peptide sequence of collagen, Gly–Pro–Hyp^14^. However, all these methods result in relatively low collagen yields or lack post-translational modifications essential for collagen function^3,15^.

An alternative approach for collagen production in mammalian cells could be mediated by the modulation of cellular pathways upregulating collagen biosynthesis. The level of various ECM components, including collagen type I, are directly related to oxygen levels and the activity of the hypoxia-inducible factor (HIF), a transcription factor that regulates the transcription of several genes, which play roles in metabolism, angiogenesis, cell survival and extracellular matrix remodeling^16–18^. Under normal oxygen levels (normoxia), conserved proline residues of HIF-1α are hydroxylated by the HIF-prolyl hydroxylase enzymes, known as prolyl hydroxylases domain (PHDs)^19–21^. Thereafter, hydroxylated HIF-1α is polyubiquitylated by the von Hippel–Lindau (VHL) tumour suppressor protein (pVHL), marking HIF-1α for proteasomal degradation^22^ **(Fig. 1a)**. However, under low oxygen levels (hypoxia), PHD becomes inactive, leading to elevated levels of the non-hydroxylated HIF-1α and the induction of a large network of HIF-target genes^16–18^, resulting in high levels of collagen type I **(Fig. 1b)**. These target genes include the collagen prolyl 4-hydroxylases and lysyl hydroxylases^23–26^. Collagen’s proline hydroxylation by collagen prolyl 4-hydroxylases is critical for stabilizing its triple helix conformation, while lysin hydroxylation by lysyl

**Figure 1.**
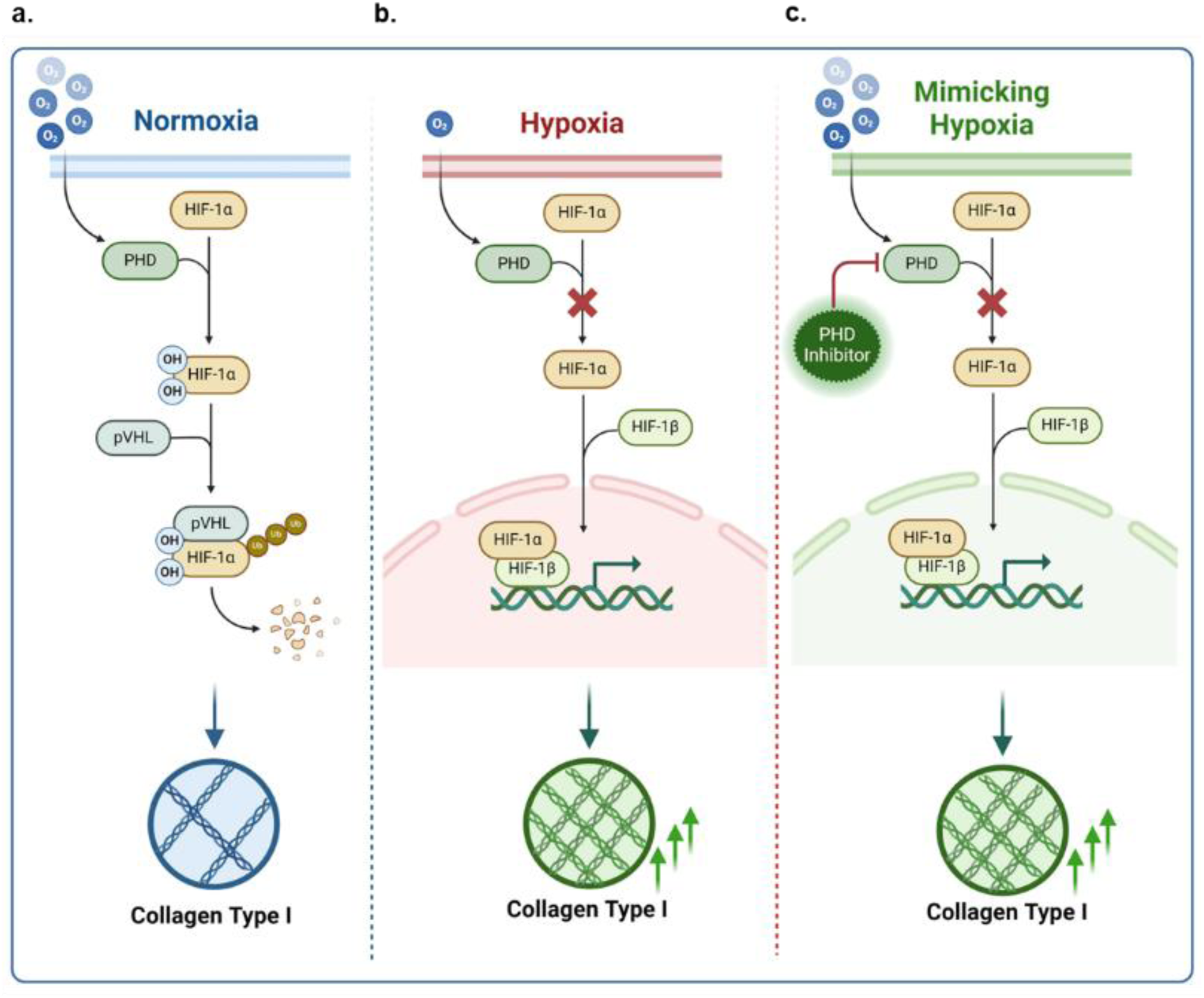
Illustration of HIF-1α activity and collagen levels under normoxia, hypoxia and hypoxia-mimicking state. **(a)** Under **normoxia**, HIF-1α is heavily hydroxylated by PHD, leading to HIF fast degradation in the proteasome. This results in basal collagen levels. **(b)** Under **hypoxia**, PHD is inactive, leading to over-activity of non-hydroxylated HIF. The HIF-1α/1β heterodimer translocates to the cell nucleus, where it transcribes hypoxia target genes, leading to over-accumulation of collagen type I. **(c) Hypoxia-mimicking state** under normal oxygen levels: small molecule PHD inhibitor prevents HIF-1α hydroxylation, leading to a similar signaling cascade as in **(b)**. Abbreviations: O2, molecular oxygen; PHD, prolyl hydroxylase domain protein; HIF-1α, Hypoxia-inducible factor 1-alpha; OH, Hydroxy group; pVHL, von Hippel–Lindau (VHL) tumour suppressor protein; Ub, Ubiquitin; HIF-1β, Hypoxia-inducible factor 1-beta. Image was created with BioRender. hydroxylases is important for collagen intermolecular cross-linking and assembly into fibrils, allowing its secretion into the extracellular space^27^.

It was shown that under hypoxia, enhanced HIF activity in human gingival fibroblasts (HGFs) and human periodontal ligament cells (HPDLs) upregulates collagen biosynthesis^28^. Moreover, we previously showed that ML228, a small molecule activator of HIF^29^, upregulates the accumulation of collagen type I in HGFs^30^. Thus, we hypothesized that mimicking a hypoxic state by directly interfering with the catalytic activity of PHD could induce HIF activation under normoxic conditions, followed by enhancement of collagen type I levels in mammalian cells **(Fig. 1c)**. In this study, we demonstrate that inhibiting PHD in mammalian cells activates the HIF transcription factor, resulting in increased collagen levels, driven by the upregulation of the key enzymes collagen prolyl 4-hydroxylases and lysyl hydroxylases. These findings suggest that mimicking hypoxia by targeting PHD represents a promising strategy for enhancing natural collagen production, offering potential therapeutic and biotechnological applications.

## Results

### The effect of PHD inhibitors on collagen type I levels

To elevate collagen levels in mammalian cells, we introduced NIH/3T3 cells with PHD inhibitors. We tested a set of 11 known small molecules, derived from 2-Oxoglutarate (2-OG), a co-substrate of HIF-1α hydroxylation^31–36^ **(Fig. 2a-b)**. Included in this set were the first-generation PHD inhibitors, N-Oxalylglycine (NOG), the first reported 2-OG mimetic molecule^37^, and Dimethyloxalylglycine (DMOG), a cell-permeable precursor of NOG^38^. Additionally, PHD inhibitors that are clinically used for treating anemia associated with chronic kidney disease (CKD)^39^ including Roxadustat^40,41^, which is approved in China and Japan for renal anemia, along with Daprodustat^42–44^, Vadadustat^45,46^, Molidustat^47^, Enarodustat^48,49^, and Desidustat^50^, which received its first approval in India. Recently, the European Medicines Agency (EMA) authorized Roxadustat to treat symptomatic anemia in CKD patients in the EU^35^. Moreover, we investigated IOX4^51^, a structurally related analog of Molidustat, FG-2216^52^, a structurally related analog of Roxadustat, and AKB-6899^53^, a structurally related compound of Vadadustat.

**Figure 2.**
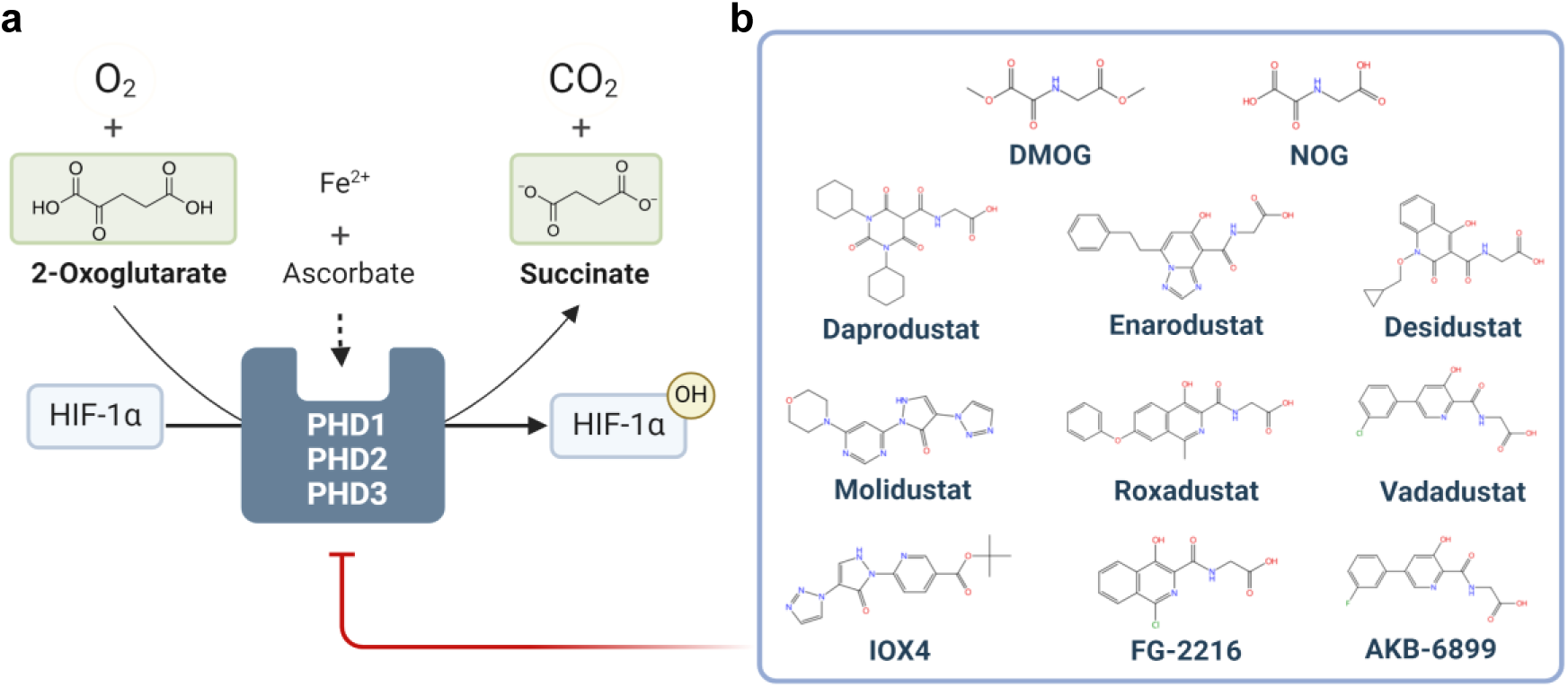
Pharmacological induction of Hypoxia-like state via PHD inhibition. **(a)** Schematic illustration of HIF-1α hydroxylation on proline residues by PHD1, PHD2, and PHD3, using molecular oxygen and 2-OG as co-substrates, and divalent iron and ascorbate as co-factors, under normoxia. The products of this enzymatic reaction are CO2, succinate and a hydroxylated HIF-1α. **(b)** Chemical structures of a set of PHD inhibitors used as competitive inhibitors of 2-OG in the current work. From left to right, top to bottom: Dimethyloxalylglycine (DMOG), N-Oxalylglycine (NOG), Daprodustat, Enarodustat, Desidustat, Molidustat and its analogues IOX4, Roxadustat and its structurally related analogue FG-2216, Vadadustat and its structurally related compound AKB-6899. The image was created with BioRender.

The NIH/3T3 embryonic mouse fibroblast cell line is known for its collagen type I production^54^. To evaluate the effect of the PHD inhibitors on collagen biosynthesis, the cells were cultured for 48 hours with various concentrations of Roxadustat, Enarodustat, Desidustat, IOX4, DMOG, NOG, Vadadustat, AKB-6899, Daprodustat, FG-2216, and Molidustat. The cells were immuno-stained for extracellular collagen type I and stained to mark the nuclei for cell quantification. Collagen levels and the number of cells were quantified via image analysis. Enhanced collagen levels are correlated with higher concentrations of Roxadustat **(Fig. 3a, j)**, Enarodustat **(Fig. 3c, k)**, and Desidustat **(Fig. 3e, m)**. Importantly, none of these compounds induced detectable cellular toxicity **(Fig. 3b, 3d, 3f)**. However, IOX4 **(Fig. 3g, n)**, DMOG **(Fig. S1a, S2b)**, NOG **(Fig. S1c, S2c)**, Vadadustat **(Fig. S1e, S2d)**, and AKB-6899 **(Fig. S1g, S2e)** did not result in a substantial increase in collagen levels. Elevated concentrations of IOX4 caused noticeable cellular toxicity **(Fig. 3h)**, whereas no such toxicity was observed with the other molecules **(Fig. S1b, S1d, S1f, and S1h)**. Daprodustat **(Fig.S1i, S2f)**, FG-2216 **(Fig. S1k, S2g)** and Molidustat **(Fig. S1m, S2h)** imparted enhanced collagen levels, though to a lower extent than Roxadustat, Desidustat and Enarodustat.

**Figure 3.**
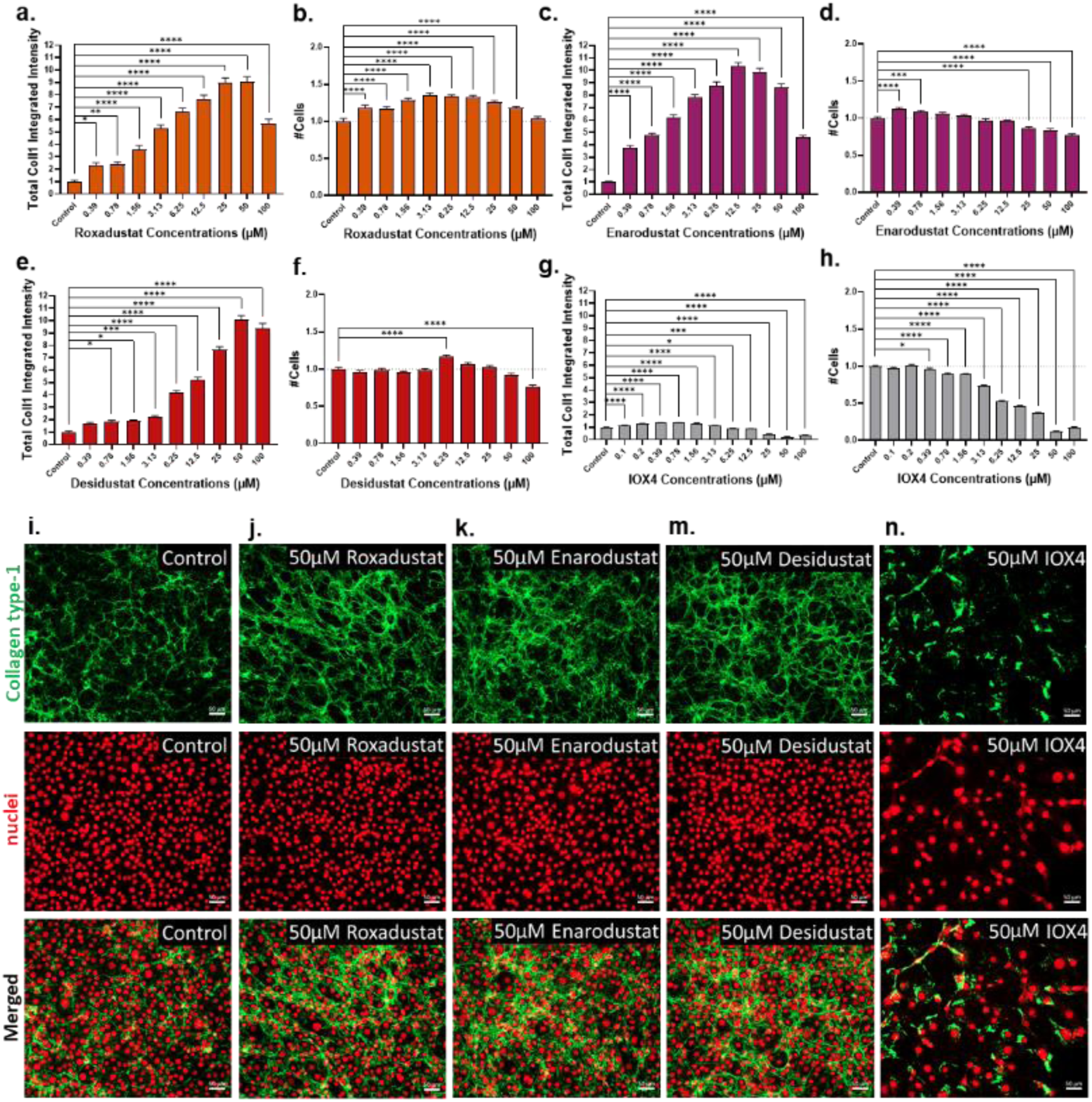
The effect of PHD inhibitors on collagen type I levels and viability in NIH/3T3 fibroblasts. Cells were cultured for 48 hours with variable concentrations of **(a-b)** Roxadustat, **(c-d)** Enarodustat, **(e-f)** Desidustat or **(g-h)** IOX4, and then labelled for collagen type I and DNA. **(a, c, e, g)** Immunofluorescence measurements of total collagen type I integrated intensity per image relative to the untreated control. **(b, d, f, h)** The number of cells relative to the untreated control. Error bars indicate standard error of the mean (SEM). Asterisks show significance following ordinary one-way ANOVA with Dunnett’s multiple comparisons test. (*) p ≤ 0.05; (**) p ≤ 0.01; (***) p ≤ 0.001; (****) p ≤ 0.0001 and (ns; not significant) p > 0.05. **(i-n)** Representative widefield images of NIH-3T3 cells 48h post-treatment. From left to right: **(i)** Untreated control, **(j)** 50 µM Roxadustat, **(k)** 50 µM Enarodustat, **(m)** 50 µM Desidustat and **(n)** 50 µM IOX4. From top to bottom: Immunostaining for collagen type I (green), staining for cell nuclei (far-red) and merged images. Scale bars, 50 µm.

No cellular toxicity was observed with the tested molecules **(Fig. S1j, S1l)**, except for Molidustat, which caused noticeable toxicity at elevated concentrations **(Fig. S1n)**.

Figure 4 summarizes the maximal fold change in collagen levels and the corresponding concentration for each molecule. Notably, Roxadustat, Desidustat, and Enarodustat yielded a significant increase in collagen type I levels of 9.0, 10.1, and 10.4 folds relative to control cells, respectively.

**Figure 4.**
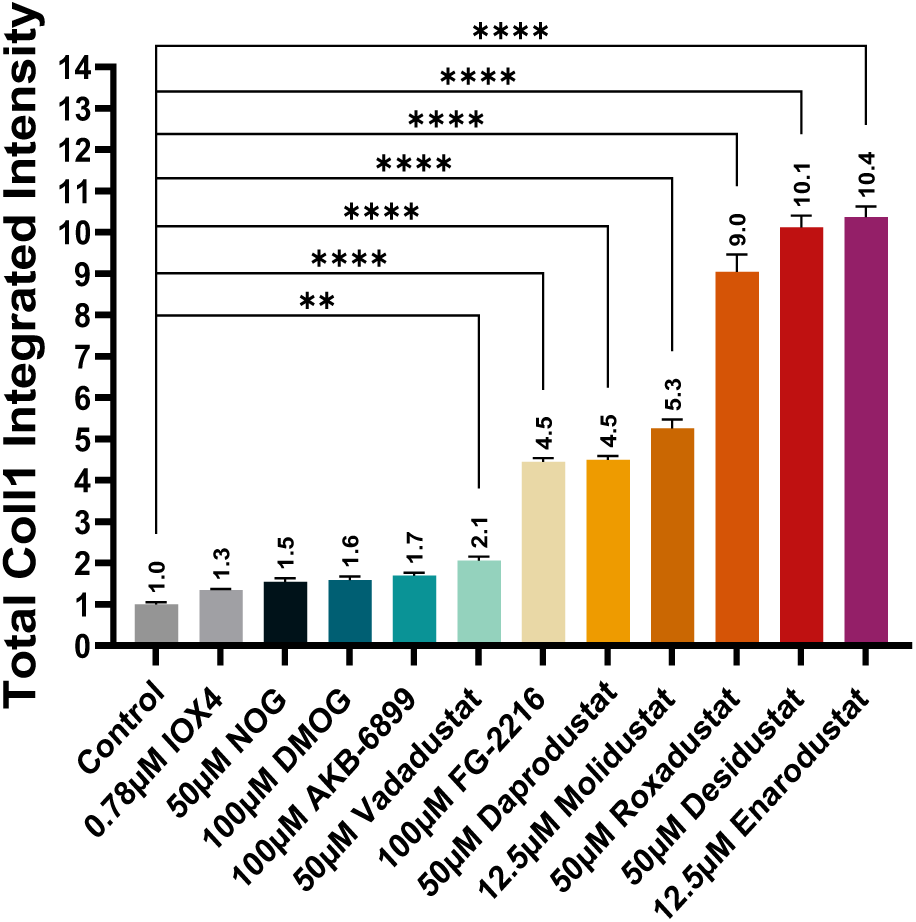
Summary of the maximum effect of PHD inhibitors on collagen type I levels. Maximum value of the total collagen type I integrated intensity relative to the untreated control for all 11 tested PHD inhibitors. Error bars indicate SEM. Asterisks show significance following ordinary one-way ANOVA with Dunnett’s multiple comparisons test. (*) p ≤ 0.05; (**) p ≤ 0.01; (***) p ≤ 0.001; (****) p ≤ 0.0001 and (ns; not significant) p > 0.05.

### Time-response and multiple-dose-treatments effect of PHD inhibitors on collagen type I levels

Next, we assessed the time-response and multiple-dose-treatment effects of Roxadustat, Enarodustat, and Desidustat, at the concentrations of 12.5 µM, 25 µM, and 50 µM respectively, in NIH/3T3 cells, selected due to their highest impact on collagen synthesis (Fig. 5). Cells were treated for 48 hours with a single application **(Fig. S3a)**, for 72 hours with two applications **(Fig. S3b)**, and for 144 hours with three applications of each of the small molecules **(Fig. S3c)**. **Figure S3** depicts the experimental design. According to collagen type I immunostaining, Roxadustat and Enarodustat yielded 26- and 29-fold increase in collagen levels, respectively, by applying the 12.5 µM treatment for 144h (Fig. 5a, c). Both molecules imparted tissue detachment from the plate at a concentration of 50 µM for 144h, limiting the ability to measure both collagen levels and the number of viable cells (Fig. 5b and 5d). The application of 25 and 50 µM Desidustat for 144h enhanced collagen levels by 23-fold (Fig. 5e), without imparting cellular toxicity (Fig. 5f). Figures 5g**-5j**, showing collagen type I immunostaining, illustrates a progressive increase in extracellular collagen over 6 days of cell growth. Notably, treatment with multiple doses of Roxadustat, Enarodustat, and Desidustat led to a significant elevation in extracellular collagen levels compared to untreated cells.

**Figure 5.**
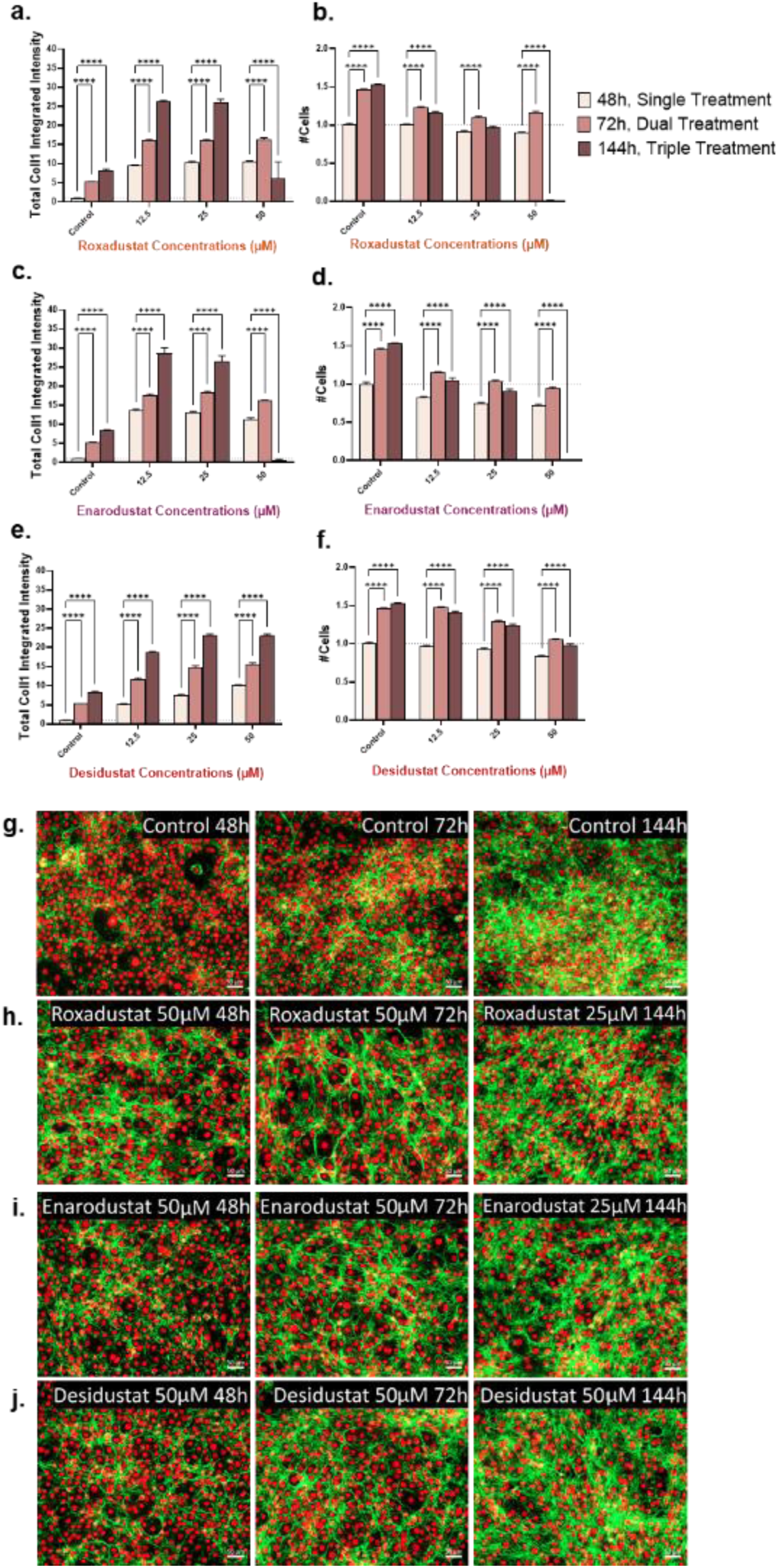
The effect of PHD inhibitors on NIH-3T3 cells at different time points. Cells were cultured for 48, 72, or 144 hours with variable concentrations of **(a-b)** Roxadustat, **(c-d)** Enarodustat, and **(e-f)** Desidustat, and then labelled for collagen type I and DNA. **(a, c, e)** Immunofluorescence measurements of total collagen type I integrated intensity per Image relative to the untreated control. **(b, d, f)** The number of cells relative to the untreated control. All data is normalized to untreated cells after 48h. Error bars indicate SEM. Asterisks show significance following ordinary one-way ANOVA with Dunnett’s multiple comparisons test. (*) p ≤ 0.05; (**) p ≤ 0.01; (***) p ≤ 0.001; (****) p ≤ 0.0001 and (ns; not significant) p > 0.05. **(g-j)** Representative widefield merged images of NIH-3T3 48h, 72h, and 144h post-treatment, immuno-stained for collagen type I (green) and stained for cell nuclei (far-red). **(g)** Untreated control, **(h)** Roxadustat, **(i)** Enarodustat, and **(j)** Desidustat at different concentrations. Scale bars, 50 µm.

### HIF-1α activation and nuclear translocation

We then studied the effect of the three best molecules, Roxadustat, Enarodustat, and Desidustat, together with Vadadustat as a reference for a molecule with low activity. We evaluated HIF-1α activation and localization within cells, focusing on both cytosolic and nuclear HIF-1α levels. NIH/3T3 cells were cultured for 48 hours with medium (Fig. 6a), supplemented with 50 µM Desidustat (Fig. 6b), 50 µM Roxadustat (Fig. 6c), 12.5 µM Enarodustat (Fig. 6d), and 50 µM Vadadustat (Fig. 6e). The cells were immune-stained for HIF-1α and stained for DNA. While HIF-1α was evenly distributed in the cytoplasm and nuclei of untreated control cells, enhanced nuclear staining was evident in the cells treated with the PHD small molecule inhibitors. Figure 6f presents an orthogonal Western blot analysis of HIF-1α protein levels in whole-cell lysates. The results indicate increased HIF-1α levels 72 hours post-treatment with Desidustat, Roxadustat, and Enarodustat compared to control cells. As expected, treatment with Vadadustat resulted in lower HIF-1α levels than the other treatments.

**Figure 6.**
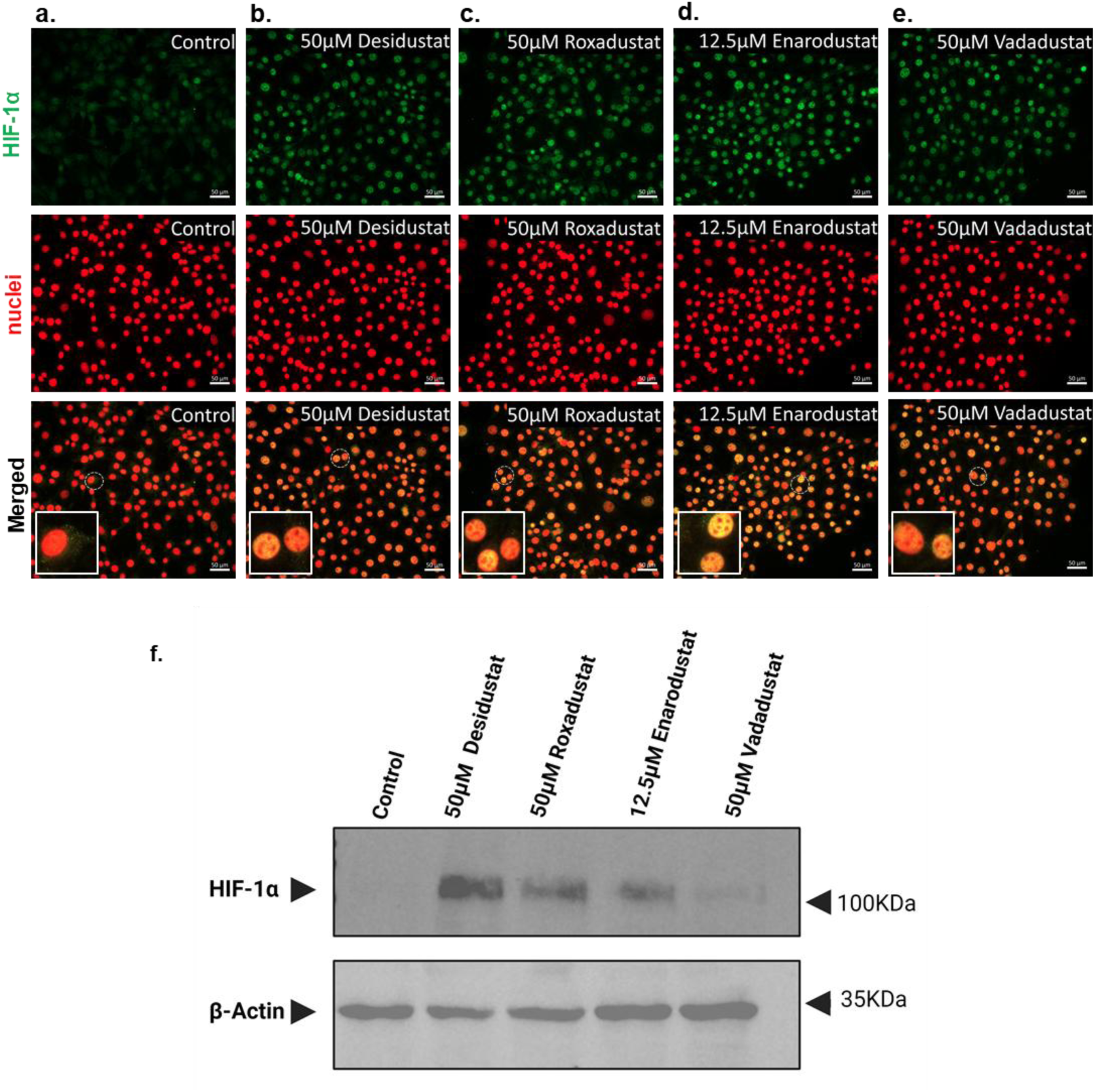
Evaluation of HIF-1α levels following treatment with PHD inhibitors. **(a-e)** Representative widefield images of NIH-3T3 cells 48h-post treatment. **(a)** Control, **(b)** 50 µM Desidustat, **(c)** 50 µM Roxadustat, **(d)** 12.5 µM Enarodustat, and **(e)** 50 µM Vadadustat. Images show cellular immunostaining for HIF-1α (green, top row), staining for DNA (red, bottom row) and merged images (bottom row). The inset in each merged image depicts a magnification of a specific region of interest (indicated by dashed lines), demonstrating the co-localization of HIF-1α with the cell nucleus. Scale bars, 50 µm. **(f)** Western blot analysis of HIF-1α levels in all treatment conditions versus control cells 72 hours post treatment. From left to right: control, 50 µM Desidustat, 50 µM Roxadustat, 12.5 µM Enarodustat or 50 µM Vadadustat. β-Actin was used as a housekeeping gene. Right arrows indicate the molecular weight of each protein.

### The effect of PHD inhibitors on the transcript levels of collagen modifying enzymes

We further tested the effect of Roxadustat and Desidustat on the transcript levels of collagen prolyl 4-hydroxylases and lysyl hydroxylases, collagen-modifying enzymes known to be regulated by HIF-1α^55–58^ (Fig. 7a). Collagen prolyl 4-hydroxylase is a α_2_β_2_ heterotetramer comprised of 2 identical α and β subunits^26,59^. The P4ha1, P4ha2 and P4hb genes encode for the α(I), α(II) and β subunits, respectively, forming collagen prolyl 4-hydroxylase I and collagen prolyl 4-hydroxylase II tetramers, respectively^23^. To evaluate the transcript levels, cells were cultured for 48h with 50 µM Roxadustat or 50 µM Desidustat, and mRNA expression level was analyzed using real-time PCR. The two molecules enhanced P4ha1 mRNA levels by ∼7-fold (Fig. 7b). Roxadustat and Desidustat enhanced P4ha2 mRNA levels by ∼15- and ∼12-fold, respectively (Fig. 7c), and both treatments enhanced P4hb mRNA levels by ∼4-fold (Fig. 7d). We also tested the mRNA levels of Plod1 and Plod2 genes, which encode the lysyl hydroxylase 1 and lysyl hydroxylase 2 isoenzymes, respectively^23,24,27^. These enzymes hydroxylate lysine residues, which serve as the only glycosylation sites on collagen molecules and function as key locations for oxidative deamination and covalent cross-linking, both essential for stabilizing collagen fibrils^24^. Roxadustat and Desidustat enhanced Plod1 by ∼7-fold (Fig. 7e), while Roxadustat and Desidustat enhanced Plod2 mRNA levels by ∼11- and ∼8-fold, respectively (Fig. 7f).

**Figure 7.**
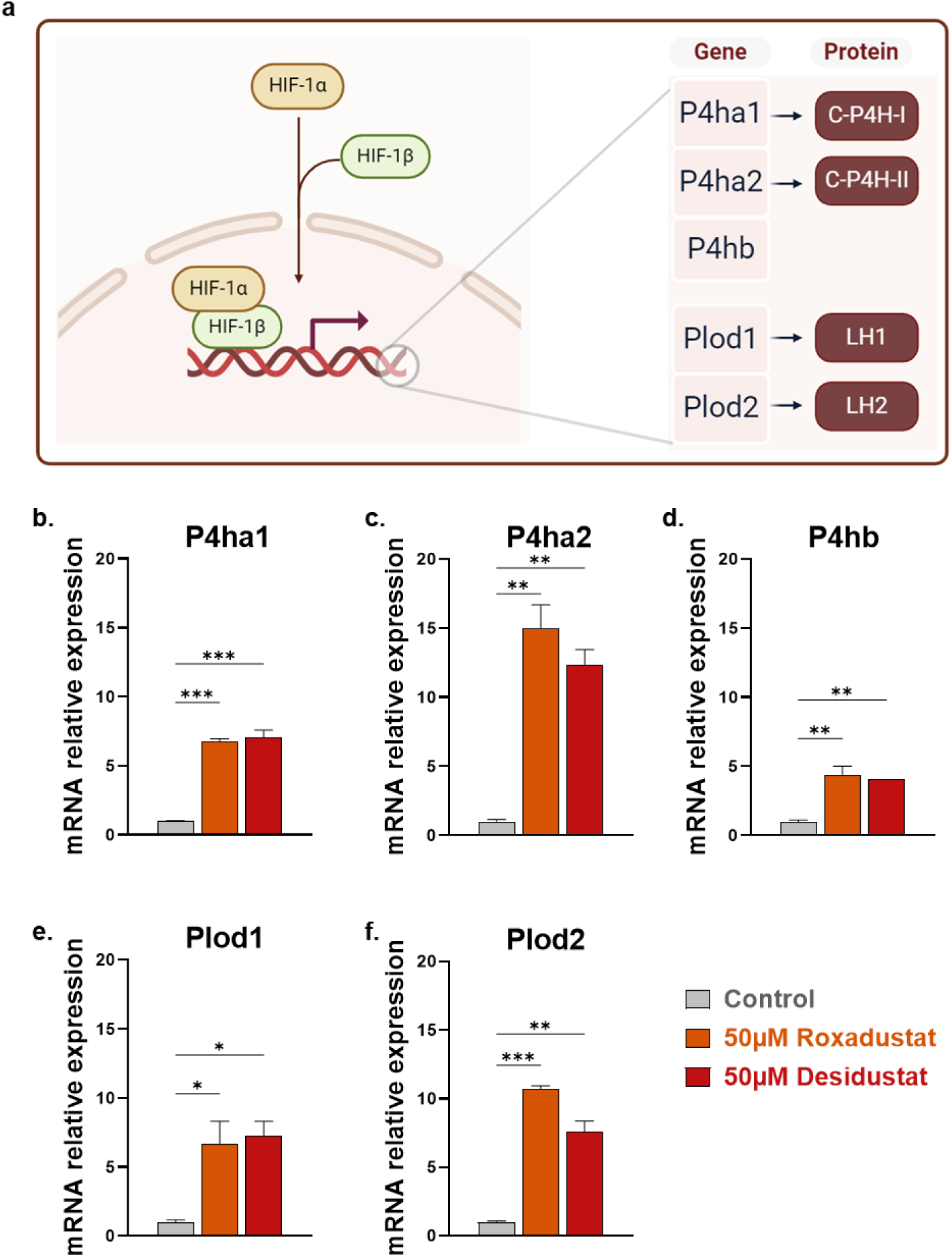
The effect of PHD inhibitors on the transcript levels of collagen modifying enzymes. **(a)** Schematic illustration of gene transcription following HIF-1α activation. Left panel: HIF-1α accumulates in the cell, forms a heterodimer with HIF-1β, and the activated HIF-1 translocate to the cell nucleus, functions as a transcription factor to hypoxia target genes. Right panel, from top to bottom: Several downstream genes to HIF-1α encode for collagen modifying enzymes: The genes P4ha1 and P4ha2 together with P4hb encode for the enzymes collagen prolyl 4-hydroxylase I and collagen prolyl 4-hydroxylase II, respectively. The genes Plod1 and Plod2 encode for the enzymes lysyl hydroxylase 1 and lysyl hydroxylase 2, respectively. **(b-f)** Cells were cultured for 48 hours with 50 µM Roxadustat or 50 µM Desidustat, and then used for real-time PCR analysis of **(b)** P4ha1, **(c)** P4ha2, **(d)** P4hb, **(e)** Plod1 and **(f)** Plod2. The 18S-rRNA housekeeping gene was used as an internal reference gene. ΔΔCT method was used for relative quantification, while fold changes were determined relative to control. Error bars indicate standard deviation (SD). Asterisks show significance following ordinary one-way ANOVA with Dunnett’s multiple comparisons test. (*) p ≤ 0.05; (**) p ≤ 0.01; (***) p ≤ 0.001; (****) p ≤ 0.0001 and (ns; not significant) p > 0.05. Abbreviations: C-P4H, Collagen prolyl 4-hydroxylase; LH, Lysyl hydroxylase; HIF-1α, Hypoxia-inducible factor 1-alpha; HIF-1β, Hypoxia-inducible factor 1-beta. Image was created with BioRender.

## Discussion

The relation between hypoxia and collagen can be used to elevate collagen synthesis in cells. In this study, we demonstrated that treating mammalian fibroblast cells with small molecule inhibitors of PHD, activates the HIF transcription factor and boosts collagen levels in the ECM. The tested 2-OG derivatives induce HIF activation under normal oxygen levels, thereby mimicking hypoxic-like conditions, leading to enhanced transcription levels of collagen prolyl 4-hydroxylase and lysyl hydroxylase genes and enhancing collagen secretion and stabilization. Our study thus integrates the relation between hypoxia and collagen, via the modulation of protein regulation cellular pathways, mimicking hypoxia under normal oxygen levels.

Collagen type I deposition and synthesis increased under hypoxic conditions in human dermal fibroblast cells^60^, corneal fibroblasts^61^, human periodontal ligament cells, and human gingival fibroblasts^28^. Additionally, studies have shown that HIF-1α regulate collagen type II synthesis in chondrocytes across several species under low oxygen levels^62–65^. Other studies suggested that HIF-1α modulates collagen secretion under hypoxic conditions by mediating hydroxylation and proper folding through collagen prolyl 4-hydroxylase as part of post-translational modifications^28,65,66^. Based on the importance of HIF in regulating collagen synthesis, we have previously shown that the application of the small molecule HIF activator ML228^29^ enhances collagen type I levels in human gingival fibroblasts^30^.

Herein, we studied the effect of 11 small molecules, all of which are derivatives of the PHD substrate 2-OG, on HIF and collagen levels in NIH/3T3 fibroblast cells. We demonstrated that these molecules effectively promote the overaccumulation of collagen type I in the ECM. Specifically, 48-hour treatment with 50 µM Roxadustat, 50 µM Desidustat or 12.5 µM Enarodustat increased collagen type I levels by 9-10-fold, compared to untreated cells, without any cytotoxicity. Additionally, treatment for 6 days with 12.5 µM Roxadustat, 12.5 µM Enarodustat, or 25 µM Desidustat boosted collagen type I levels by ∼26-fold, 29-fold, and 23-fold, respectively, compared to untreated cells.

HIF-1α serves as a key regulator of the cellular response to hypoxia, orchestrating the transcription of genes involved in adaptive processes, including those critical for collagen biosynthesis and its post-translational modifications^55–58^. In our study, 48 and 72-hour treatments with 50 µM Desidustat, 50 µM Roxadustat, or 12.5 µM Enarodustat resulted in a significant elevation of HIF-1α protein levels, as confirmed by immunostaining and Western blot analysis, respectively, likely due to reduced proteasomal degradation of HIF-1α. Additionally, 48 hours post-treatment with 50 µM Roxadustat or 50 µM Desidustat, we observed increased transcription of genes encoding collagen prolyl 4-hydroxylases (P4ha1, P4ha2, P4hb) and lysyl hydroxylases (Plod1, Plod2), enzymes that play a critical role in stabilizing the collagen triple helix and promoting fibril assembly. These results are consistent with the mechanistic link between PHD inhibition, HIF-1α stabilization, and the overaccumulation of collagen, possibly driven by enhanced collagen hydroxylation and maturation.

Our study shows a novel approach for boosting natural collagen production in mammalian cells by specific modulation of protein signaling cascades, and specifically by imparting a hypoxia-like condition in the cells. However, we also note a few limitations of the small molecules we tested. Despite the overall positive effect on collagen levels, these small molecules may lack specificity. The collagen prolyl 4-hydroxylases and lysyl hydroxylases, like PHDs, belong to the 2-OG-dependent oxygenase superfamily^23,24,27,67^. It is, therefore, possible that the small molecule inhibitors derived from 2-OG inhibit collagen prolyl 4-hydroxylases and lysyl hydroxylases, potentially limiting further increases in collagen type I levels. Such dual effect could be solved by using orthogonal methods such as siRNA that will specifically inhibit PHD transcripts or by developing other modulators that will interfere with PHD/HIF protein-protein interaction rather than binding to PHD catalytic site.

In addition, multiple cellular bottlenecks might hamper collagen biosynthesis. Thus, future efforts should explore the integration of multiple synergistic strategies. For instance, the incorporation of ascorbic acid (vitamin C), an important collagen prolyl 4-hydroxylase and lysyl hydroxylase co-factor^23,24,27,67^, could further enhance collagen biosynthesis. Furthermore, exploring the inhibition of key regulatory points in the HIF pathway, such as the VHL ^68^, could lead to additional enhancement of collagen levels. Additional orthogonal approaches, such as enhancing the levels of collagen prolyl 4-hydroxylase and lysyl hydroxylase enzymes, could boost the collagen biosynthesis rate. Given collagen’s pivotal role across diverse fields, employing 2-OG derivatives to inhibit PHD and the application of alternative schemes for hyper-activation of the HIF pathway represents a promising approach to achieve high-yield expression of natural, cell-derived collagen.

## Materials and Methods

### Small Molecules

All 11 small molecules were obtained from the Aldrich Market Select Small Molecule Screening Library Service, with a purity >95%, and dissolved in DMSO. DMOG (Dimethyloxalylglycine; CAS: 89464-63-1), NOG (N-Oxalylglycine; CAS: 5262-39-5) and FG-2216 (IOX 3; CAS: 223387-75-5) were dissolved to a stock solution of 100 mM. Roxadustat (FG 4592; CAS: 808118-40-3), Enarodustat (JTZ 951; CAS: 1262132-81-9), Desidustat (ZYAN 1; CAS: 1616690-16-4), IOX4 (CAS: 1154097-71-8), Vadadustat (AKB 6548; CAS: 1000025-07-9) and AKB-6899 (CAS: 1007377-55-0) were dissolved to a stock solution of 50 mM. Daprodustat (GSK 1278863; CAS: 960539-70-2) and Molidustat (BAY 85-3934; CAS: 1154028-82-6) were dissolved to a stock solution of 25 mM. Desidustat, Daprodustat, and Molidustat underwent heat-sonication before use.

### Cell Culture and Treatment with Small Molecules for 48 hours

NIH/3T3 embryonic mouse fibroblast cell line (ATCC, CRL-1658) was cultured in Dulbecco’s Modified Eagle’s Medium (DMEM, high glucose, no glutamine) (Gibco) supplemented with 10% Fetal Bovine Serum (Gibco), 1% L-Glutamine (200 mM) (Sartorious), and 1% Penicillin-Streptomycin (Sigma-Aldrich). Cells were maintained at 37°C in a humidified incubator with 5% CO₂. Twenty-four hours after cell seeding, the cells were treated for 48 hours with a serial dilution of each of the 11 small molecules or with DMSO as a control. The final DMSO concentration in the culture medium was matched to the small molecule treatments: 0.1% for DMOG, NOG, and FG-2216; 0.2% for Roxadustat, Enarodustat, Desidustat, IOX4, Vadadustat, AKB-6899, and Daprodustat; and 0.4% for Molidustat.

### Collagen type I immunofluorescence and DNA staining

NIH/3T3 cells were seeded at a concentration of 5,000 cells per well in 96-well glass bottom plates (P96-1.5H-N, Cellvis). After 24 hours, cells were treated for 48 hours with media supplemented with various PHD molecules or with DMSO as a control. At the end of the 48-hour treatment period, cells were fixed for 10 minutes using 8% paraformaldehyde (PFA; Electron Microscopy Sciences) dissolved in phosphate-buffered saline (PBS; Invitrogen) to a final concentration of 4% PFA, without any washing to avoid cell detachment from the glass. The cells were then briefly washed four times with ice-cold PBS. Next, a blocking solution containing 1.5% Bovine Serum Albumin (Sigma Aldrich) dissolved in PBS was applied for 30 minutes, followed by an overnight incubation at 4°C with Rabbit recombinant monoclonal anti-mouse Collagen type I antibody [EPR24331-53] (Abcam, ab270993), dissolved in the blocking solution (1:1000 dilution). The following day, cells were washed three times with PBS (5 minutes each), followed by immunostaining with Alexa Fluor 488-conjugated goat anti-rabbit IgG H&L (Abcam, ab150077) dissolved in blocking solution (1:1000 dilution). Cells were washed again three times with PBS (5 minutes each), then stained for 30 minutes with DRAQ5™, a cell-permeable far-red fluorescent DNA dye (Abcam, ab108410) dissolved in PBS (1:1000 dilution), and washed once more with PBS.

### Confocal microscopy

Images were acquired using a ZEISS LSM 900 microscope in widefield detection mode. Fluorescence readouts were obtained at excitation/emission of 493/517 nm (green) and 647/683 nm (far-red) using Objective Plan-Apochromat 20x/0.8 M27 for collagen detection or Objective LD LCI Plan-Apochromat 25x/0.8 Imm Corr DIC M27 for HIF-1α detection.

### Image analysis

Image acquisition and quantification were performed using the Incucyte SX5 (Sartorius) software, version 2023A. A standard scan was performed for 9 images per well using objective 20x, 2 optical modules (green at excitation/emission of 453–485 nm/494–533 nm in 300 msec acquisition time, and near-IR at excitation/emission of 648-674 nm/685-756 nm in 400 msec acquisition time) following immunofluorescence protocol. For image analysis, 2-6 replicates per condition were used. Using the Basic Analyzer mode, “Total Collagen type I Integrated Intensity Per Image (Green calibrated unit (GCU) x µm²/Image)” was acquired using the green optical module, providing the total sum of the collagen type I objects’ fluorescence intensity per image. “# Cells (Per image)” was acquired using the near-IR optical module by counting the number of nuclei per image.

### Cell culture and treatment with small molecules at different time points

For the time points experiment, cells were re-supplemented for 48, 72, or 144 hours (see experimental design in Fig. S3) with either 12.5 µM, 25 µM, or 50 µM of Roxadustat, Enarodustat, and Desidustat, or with 0.1% DMSO as a control.

### HIF-1α immunofluorescence and DNA staining

NIH/3T3 cells were seeded at a concentration of 5,000 cells per well in 96-well glass bottom plates (P96-1.5H-N, Cellvis). After 24 hours, cells were treated for 48 hours with either 50 µM Desidustat, 50 µM Roxadustat, 12.5 µM Enarodustat, 50 µM Vadadustat, or 0.1% DMSO as a control.

At the end of the 48-hour treatment period, cells were fixed for 10 minutes with 4% paraformaldehyde (PFA, Electron Microscopy Sciences) dissolved in phosphate-buffered saline (PBS, Invitrogen) to a final concentration of 4% PFA. Cells were briefly washed three times with PBS to remove the PFA and kept overnight at 4°C. The following day, a blocking solution containing 0.1% Triton X-100 (X100, Sigma-Aldrich) and 5% BSA (Bovine Serum Albumin) (Sigma Aldrich) dissolved in PBS was applied for 1 hour, followed by 72 hours incubation at 4°C with Rabbit recombinant monoclonal anti-mouse HIF-1 alpha antibody [EPR16897] (abcam, ab179483), dissolved in 0.5% BSA in PBS (1:500 dilution). The following day, cells were washed 3 times with PBS, followed by immunostaining with Alexa Fluor 488-conjugated goat anti-rabbit IgG H&L (Abcam, ab150077), dissolved in 0.5% BSA in PBS (1:1000 dilution). Cells were washed again for 3 times with PBS, stained for 5 minutes with DRAQ5™ (ab108410, Abcam) for nuclei staining dissolved in PBS (1:1000 dilution), and washed three times with PBS.

### HIF-1α western blot analysis

NIH/3T3 cells were seeded at a concentration of 15,000 cells/cm² in 100 mm plates. After 24 hours, the cells were treated for 72 hours with either 50 µM Desidustat, 50 µM Roxadustat, 12.5 µM Enarodustat, 50 µM Vadadustat, or 0.1% DMSO as a control. At the end of the 72-hour treatment period, the cells were trypsinized, washed with PBS, and centrifuged. A total of 1x10^6^ cells were collected from each sample. The PBS was removed, and the cells were stored at -80°C until further analysis. Cells were sonicated on ice and lysed in a buffer containing cOmplete™, Mini, EDTA-free Protease Inhibitor Cocktail (#11836170001, Merck), 10 mM Phenylmethylsulfonyl fluoride (PMSF-RO, #10837091001, Merck), 1 mM Sodium orthovanadate (S6508, Merck) and 1% sodium dodecyl sulfate (BP166, Fisher scientific). Total protein was determined using Pierce BCA protein assay kit (23225, Thermo scientific) and 15 µg of total protein was loaded in each lane of a 10% SDS polyacrylamide gel, electrophoresed at 120V for 20 minutes and then at 160V for another 40 minutes. Then, protein was wet transferred to a nitrocellulose membrane at 80 mA for 60 minutes. The membrane was blocked for 2h using 5% BSA (Sigma Aldrich) in TBS-T (Tris Buffered saline) containing 0.5% Tween 20, 150 mM NaCl and 50 mM Tris-HCl at pH 7.6 on a shaker. The membrane was cut in between the 60-75 KDa ladder mark. The upper part of the membrane was incubated with Rabbit recombinant monoclonal anti-mouse HIF-1 alpha antibody [EPR16897] (ab179483, abcam) at a 1:500 dilution in 0.5% BSA in TBS-T, and the bottom part of the membrane was incubated with Mouse Monoclonal anti mouse beta Actin antibody (ab8226, abcam), at a 1:1000 dilution in 0.5% BSA in TBS-T at 4°c overnight with shaking. Membranes were washed 3 times with TBS-T with shaking (5 minutes each). Then, the membranes were incubated with Goat Anti-Rabbit IgG Antibody, HRP-conjugate (12-348, Merck) at room temperature with shaking for 1 hour, and then washed 3 times with TBS-T with shaking (5 minutes each). Next, Electrochemiluminescence (ECL) assay was performed using 2 ml HRP substrate (WBLUR0500, Millipore) for 3 minutes. Images were acquired using ChemiDoc imager (Bio-Rad, CA).

### Real-time polymerase chain reaction (RT-PCR)

NIH/3T3 cells were seeded at a concentration of 150,000 cells per well in a 6-well plate (Corning). After 24 hours, the cells were treated for 48 hours with either 50 µM Roxadustat, 50 µM Desidustat, or 0.1% DMSO as a control. At the end of the 48-hour treatment period, up to 3x10^6^ cells were collected and frozen at -80℃ for further assessment. Total RNA was isolated using NZY Total RNA Isolation kit (nzytech, MB13402) according to the manufacturer’s instructions. cDNA was synthesized from total RNA using AzuraQuant™ cDNA Synthesis Kit (Azura Genomics, AZ-1996) according to the manufacturer’s instructions. Real-time PCR was performed using AzuraViewTM GreenFast qPCR Blue Mix LR (Azura Genomics,AZ-2305) in a 96 well PCR plate (Lifegene), with 10 ng cDNA in a 20 μl total PCR reaction, using QuantStudio™ 5 Real-Time PCR System (Thermo Fisher Scientific Inc.). Primers were ordered from IDT (Integrated DNA Technologies, see Table 1). mRNA expression was analyzed based on 2-3 biological replicates each with 3 technical repeats using QuantStudio™ Design & Analysis Software (Thermo Fisher Scientific Inc.). The 18S-rRNA housekeeping gene was used as an internal reference gene. The ΔΔCT method was used for relative quantification, and fold changes were determined relative to the control.

**Table 1:**
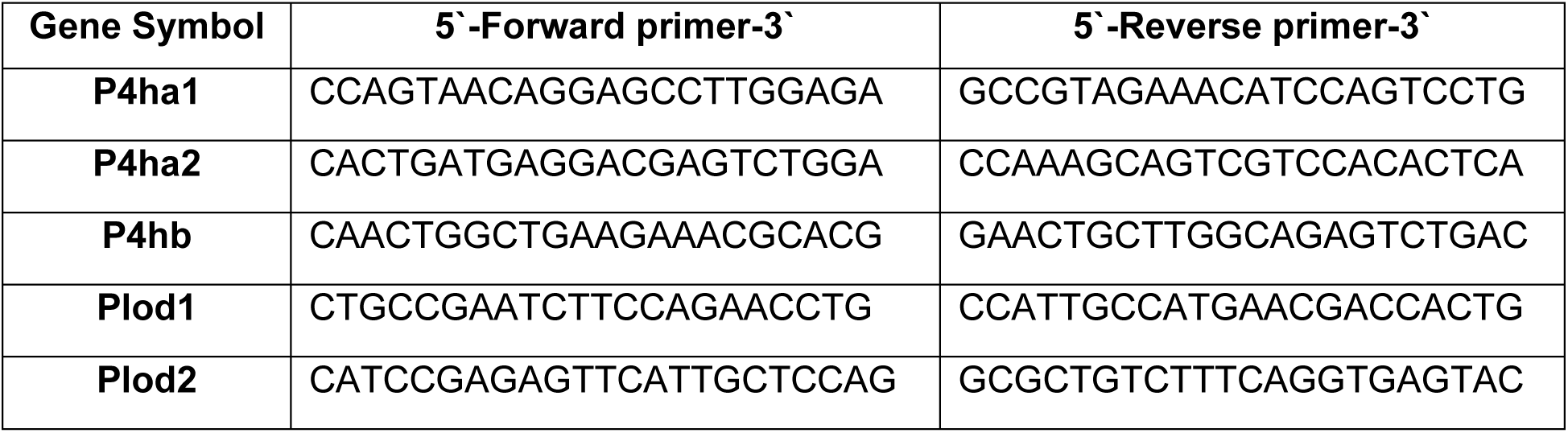
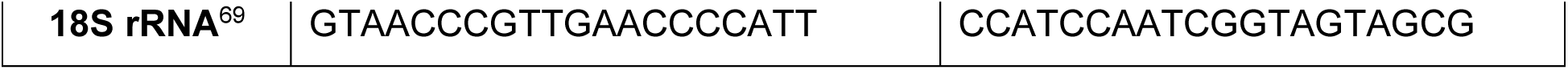
Primers used to for real-time PCR analysis.

## Statistical analysis

Multiple comparisons were performed using ordinary one-way ANOVA followed by Dunnett’s post-hoc statistical hypothesis for comparing each condition mean to the control mean. Statistical analysis was performed using GraphPad Prism version 10.0.0 for Windows, GraphPad Software, Boston, Massachusetts USA, worldwideweb(dot) graphpad(dot)com. For statistical analysis, significance was set as (*) p ≤ 0.05; (**) p ≤ 0.01; (***) p ≤ 0.001; (****) p ≤ 0.0001 and (ns; not significant) p > 0.05.

## Acknowledgements

This work was partially supported by the European Research Council (ERC), under the European Union’s Horizon 2020 research and innovation program (grant agreement no. 948102) (L. A.-A.), the Israel Science Foundation (ISF grant: 2422/24) (L. A.-A.), and the Ministry of Innovation, Science and Technology (grant 0005749) (M.G). Lucia Adriana Lifshits is grateful for Ph.D. scholarship support from the ADAMA Center at Tel Aviv University. The authors would like to acknowledge the assistance of the Research Infrastructure Core Facilities at the Faculty of Medical & Health Sciences of Tel-Aviv University. Special thanks to Dr. Yael Zilbershtein, a staff scientist, for her expert help with the Incucyte SX5 scanning and analysis. The authors acknowledge the Chaoul Center for Nanoscale Systems at Tel Aviv University for the use of instruments and staff assistance. Our gratitude extends to Dr. Yaara Oren and Rinat Semyatich for their support with the Incucyte SX5. We are thankful to Vania Altobelli for her contributions to the design of the figures and to Shir Givon for her support throughout the project. Additionally, we thank Sigal Rencus-Lazar for her assistance with editing. We thank the members of the Adler-Abramovich group, Gal group, and Arrakis Bio for their helpful discussions.

## Ethics declarations Competing interests

L.A.A. M.G. and D.R. are the co-founders of Arrakis Bio Ltd., developing animal-free collagen. All other authors declare no conflicts of interest.

