## Supplementary Information for "Enhancing Collagen Biosynthesis in Mammalian Cells Through Hypoxia-Mimetic Prolyl Hydroxylase Inhibition"

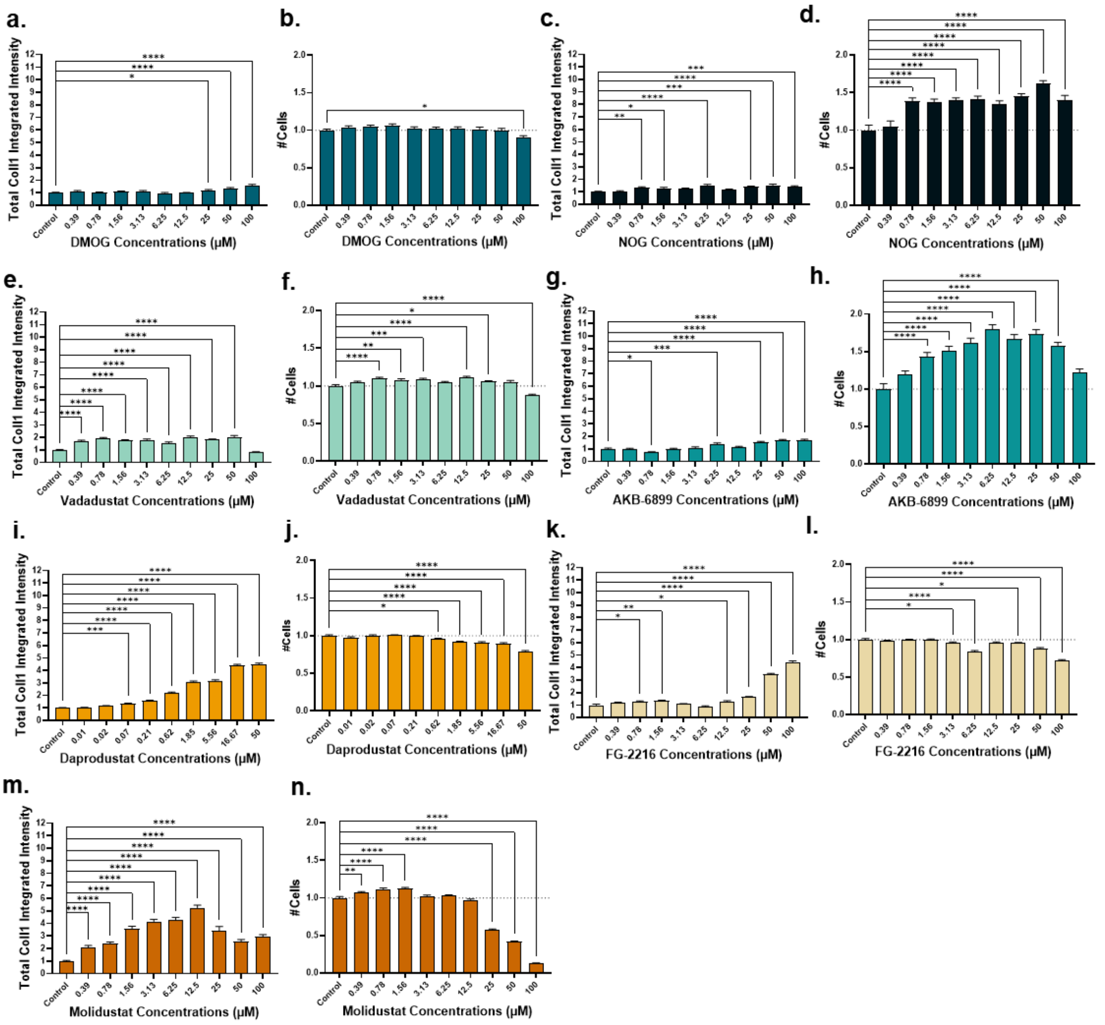


**Figure S1. The effect of PHD inhibitors on collagen type I levels and viability in NIH/3T3 fibroblasts:** Cells were cultured for 48 hours with variable concentrations of **(a-b)** DMOG, **(c-d)** NOG, **(e-f)** Vadadustat, **(g-h)** AKB-6899, **(i-j)** Daprodustat **(k-l)** FG-2216 or **(m-n)** Molidustat, and then labeled for collagen type I and DNA. **(a, c, e, g, i, k, m)** Immunofluorescence measurements of total collagen type I integrated intensity per image relative to the untreated control. **(b, d, f, h, j, l, n)** The number of cells relative to the untreated control. Error bars indicate SEM. Asterisks show significance following ordinary one-way ANOVA with Dunnett's multiple comparisons test. (*) $p \le 0.05$; (**) $p \le 0.01$; (***) $p \le 0.001$; (****) $p \le 0.0001$ and (ns; not significant) $p > 0.05$.

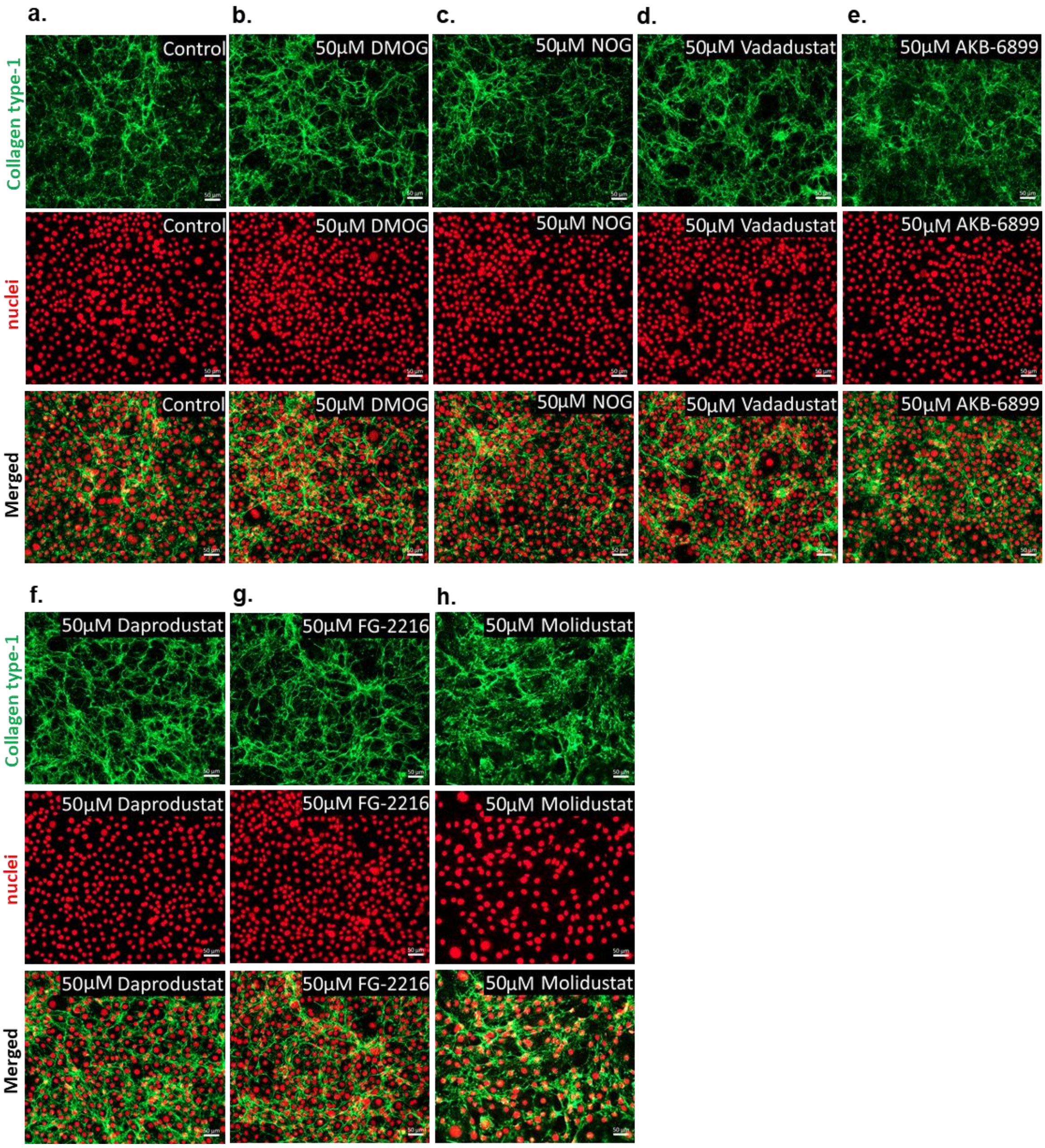


**Figure S2. The effect of PHD inhibitors on collagen type I levels:** Representative Widefield images of NIH-3T3 48h-post treatment. Upper panel, from left to right: **(a)** control, **(b)** 50 µM DMOG, **(c)** 50 µM NOG, **(d)** 50 µM Vadadustat **(e)** 50 µM AKB-6899. Lower panel, from left to right: **(f)** 50 µM Daprodustat, **(g)** 50 µM FG-2216, and **(h)** 50 µM Molidustat. Samples were then labeled for collagen type I and DNA. From top to bottom: Immunostaining for collagen type I (green), staining for cell nuclei (far-red), and merged images. Scale bars, 50 µm.

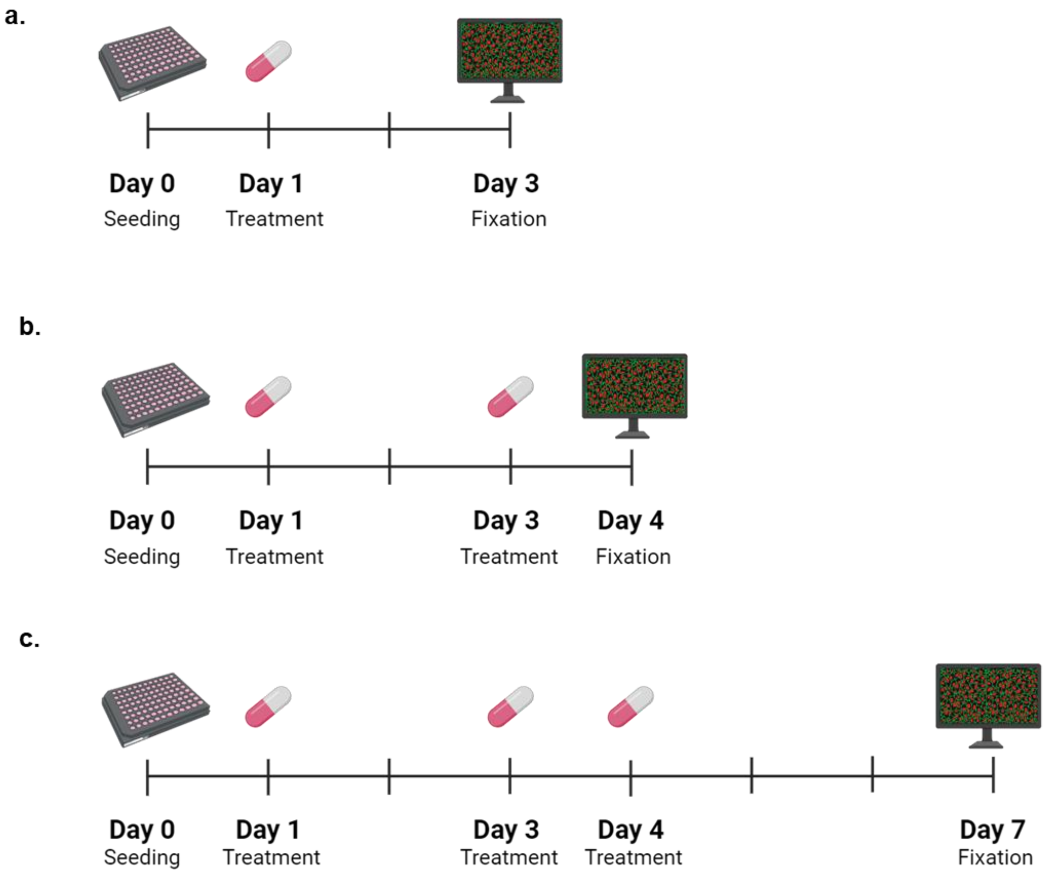


**Figure S3. The effect of multiple PHD inhibitors treatment on collagen type I levels – experimental design:** NIH-3T3 cells were cultured for different durations with 12.5 µM, 25 µM, or 50 µM of Roxadustat, Enarodustat, or Desidustat, and then labeled for collagen type I and DNA. **(a)** In plate 1, cells were treated once for 48 hours and then fixed for immunofluorescence analysis. **(b)** In plate 2, cells were treated twice over a 72-hour period, with the molecules reapplied after 48 hours, followed by fixation for immunofluorescence analysis. **(c)** In plate 3, cells were treated three times over a 144-hour period, with the molecules reapplied after 48 hours and 72 hours, followed by fixation for immunofluorescence analysis. Image created with BioRender.com.
